# Towards Unraveling Biomolecular Conformational Landscapes with a Generative Foundation Model

**DOI:** 10.1101/2025.05.01.651643

**Authors:** The OpenComplex Team, Qiwei Ye

## Abstract

Recent advances in deep learning have revolutionized biomolecular structure prediction, yet a significant challenge remains: biological function emerges from dynamic equilibria across multiple conformational states rather than a single molecular structure. Here, we present OpenComplex2 (OC2), a generative model that efficiently sample biomolecular conformational ensembles. OC2 integrates a unified graph representation capturing diverse molecular inputs with diffusion-based sampling through our FloydNetwork architecture, which operates simultaneously across atomic, residue, and motif hierarchies. This multi-level approach balances fine-grained precision with computational efficiency, enabling conformational landscape sampling in hours rather than the weeks required by traditional simulations. Through comprehensive evaluation, we demonstrate that OC2 accurately reproduces experimentally determined protein and RNA ensembles, generates conformational distributions comparable to millisecond-scale molecular dynamics simulations, and captures functionally relevant transitions in important biological scenarios including allostery, induced-fit mechanisms, and cryptic pocket formation. OC2 also effectively models small molecule conformations with high validity, identifies multiple binding sites, and maintains competitive accuracy on structure prediction benchmarks, while scaling to ultra-large molecular assemblies exceeding 15,000 residues. By bridging static structure prediction and dynamic ensemble characterization, OC2 advances our understanding of how biomolecular dynamics give rise to biological function.

## 1 Unifying Structure Prediction and Conformational Ensemble Modeling Using OpenComplex2

Proteins are inherently dynamic under physiological conditions, and their functions are typically determined by an equilibrium distribution across multiple conformational states [1]. When a drug molecule binds to its target, proteins often undergo dramatic conformational changes that directly impact therapeutic efficacy. For instance, HIV-1 protease—a critical drug target—exhibits significant structural rearrangements upon inhibitor binding that determine treatment success [2]. This intrinsic structural heterogeneity underpins a wide range of biological processes, including enzyme catalysis, molecular recognition, allosteric regulation, and signal transduction [3].

Experimental structure-determination techniques such as X-ray crystallography [4] and cryo-electron microscopy (cryo-EM) [5], are designed to resolve high-resolution structures of highly stable states of biomolecules, while other techniques, like Nuclear magnetic resonance (NMR) spectroscopy [6], can offer dynamic insights but become challenging for large systems. Current approaches for recovering equilibrium distributions still depend on classical simulations that remain computationally prohibitive for large assemblies [7]. Despite advances in specialized hardware (*e*.*g*., Anton [8, 9]) and enhanced sampling methods [10–12], achieving biologically-meaningful timescales (typically hundreds of milliseconds) for moderate-sized proteins still demands thousands of computing hours [7].

While deep learning models have demonstrated remarkable success in predicting static biomolecular structures [13–19], they remain fundamentally limited to single-state prediction and lack dynamic mechanistic insights. Recent advancements have combined AlphaFold2 [13] with Multiple Sequence Alignment (MSA) manipulation [20, 21] or generative modeling approaches [22, 23] to sample conformational diversity. However, these methods still struggle to quantitatively reproduce thermodynamic distributions and achieve atomic-scale agreement with experimental observables. Thus, a comprehensive understanding of biomolecular function requires methods that can both accurately predict stable conformations and efficiently sample equilibrium ensembles across biologically relevant timescales. We present OpenComplex2 (OC2), a generative foundation model designed to achieve these dual objectives for diverse biomolecular systems.

OC2 utilizes a score-based diffusion framework [24] to gradually transform a simple distribution (*e*.*g*., a Gaussian) into the complex equilibrium distribution of molecular systems (Fig. 1a). By enabling independent sampling from the learned distribution, OC2 explores conformational space orders of magnitude faster than traditional molecular dynamics simulations. The diffusion process is carried out by FloyedNetwork, a graph neural network that processes system-specific descriptors (*e*.*g*., protein sequences, ligand SMILES) while hierarchically integrating molecular information across scales (*e*.*g*., atom, residue, and motif) to yield complete structural dynamics representations. To enable joint learning of diverse biomolecular entities, OC2 employs a unified graph representation that standardizes heterogeneous molecular inputs into a consistent format (Fig. 1b). Notably, to extend OC2’s applicability to ultra-large biomolecular complexes, we introduce *Isomorphic Subgraph Merge* (ISM), a technique that identifies and processes symmetry-related subgraphs collectively, thereby eliminating redundant computation without sacrificing structural resolution.

**Fig. 1.**
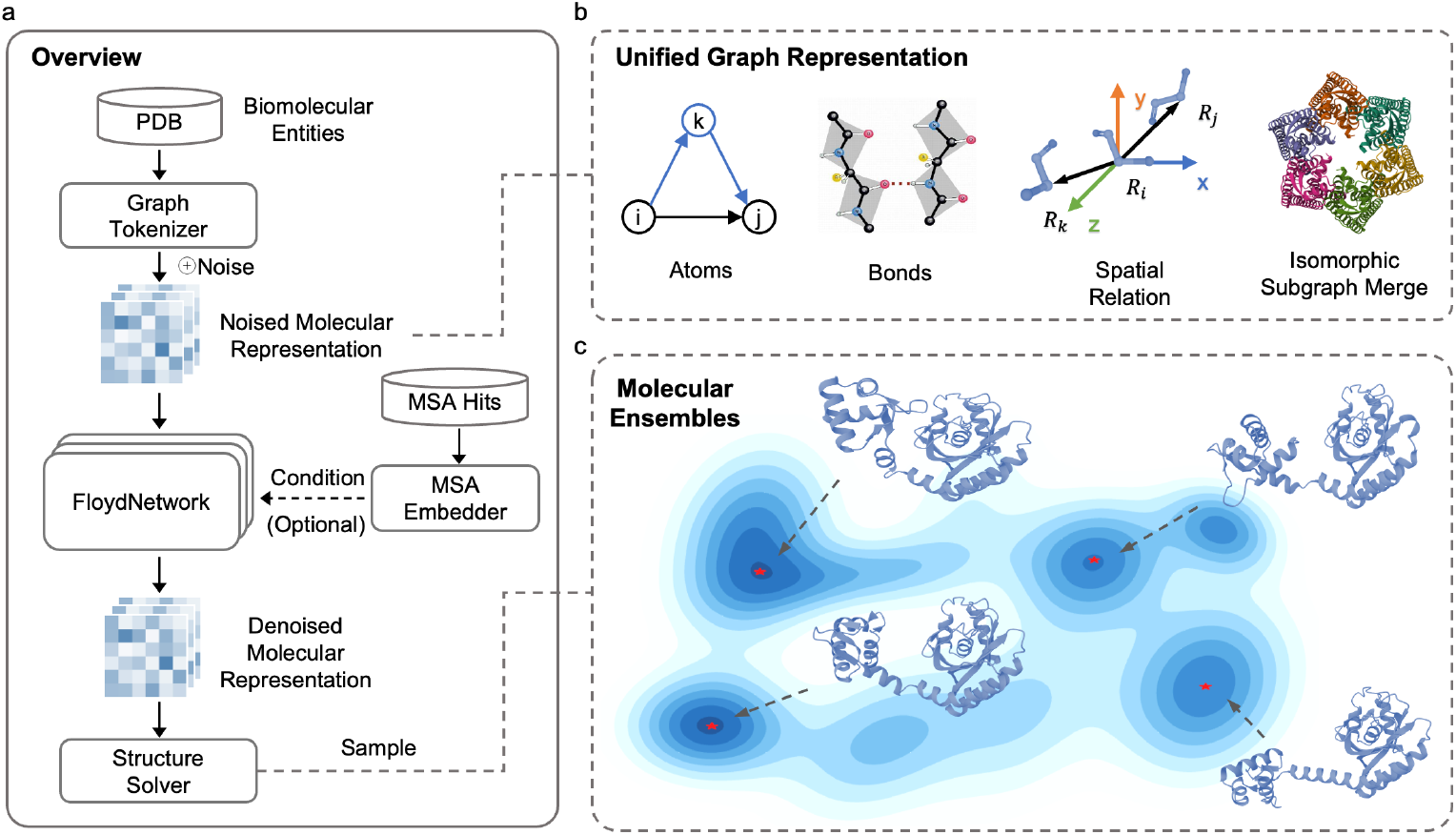
Predicting conformational landscape with the OC2 framework. **a**, Conditioned on input biomolecular entities (*e*.*g*., polymer sequences and ligand SMILES), OC2 employs a diffusion-based generative model to produce structures. This diffusion process is implemented via FloydNetwork—a graph neural network that captures interactions across atomic, residue, and motif scales. **b**, Molecular entities are transformed into a *unified* graph representation via a graph tokenizer, with the ISM method enabling efficient handling of symmetric large complexes. **c**, Through innovations in data representation and training, OC2 efficiently explores conformational landscapes.

For training, OC2 primarily utilizes biomolecular structures from the Protein Data Bank (PDB) with a cutoff date of 30 September 2021. These structures were rigorously curated and processed to extract accurate molecular representations with complete topological details. To enhance biological fidelity and capture physiologically relevant molecular interactions, we preserved all native biological partners in complexes during training, allowing OC2 to model environmental effects that influence conformational distributions. Full details of the dataset preparation are provided in Supplementary A. By incorporating these design principles, OC2 effectively bridges static structure prediction and dynamic ensemble modeling, enabling efficient generation of thermodynamic conformational ensembles without compromising structural accuracy (Fig. 1c).

We evaluate OC2’s performance across two key dimensions: **(i)** equilibrium ensemble sampling and **(ii)** high-accuracy structure prediction. For ensemble modeling, OC2 successfully: (a) reproduces NMR-derived protein and RNA conformational distributions (Sec. 2.1.1), (b) achieves sampling comparable to millisecond molecular dynamics simulations (Sec. 2.1.2), and (c) captures functionally relevant transitions including induced-fit binding and cryptic pocket formation (Sec. 2.1.3). Notably, OC2 can facilitate drug discovery by efficiently exploring diverse protein-ligand binding poses and identifying multiple binding sites (Sec. 2.3.2). For structure prediction, OC2 demonstrates: (a) state-of-the-art (SOTA) accuracy on standard benchmarks (Sec. 2.2.1), (b) scalable modeling of ultra-large assemblies (*>*15,000 residues; Sec. 2.2.2), and (c) precise stereochemical modeling of small molecules (Sec. 2.3.1). These capabilities establish OC2 as a transformative tool for statistical mechanistic studies, with broad applications in drug discovery, protein design, and biochemical mechanism elucidation.

## 2 Results

Here, we present the evaluation results of OC2 on two challenging tasks, conformational distribution prediction (Sec. 2.1) and biomolecular structure prediction (Sec. 2.2). Considering the importance of small molecules in biological processes and the drug discovery loop, we further highlight OC2’s capability in modeling small molecule conformations in complex protein-ligand environments, including accurate pose prediction and identification of multiple binding sites (Sec. 2.3). Implementation details are provided in Supplementary B.

### 2.1 Conformational Distribution Prediction

#### 2.1.1 Reproducing thermodynamic ensembles

OC2 effectively sample diverse conformations from input polymer sequences and ligand SMILES. In Fig. 2a-b, we demonstrate its ability to capture thermodynamic distributions using a set of NMR spectroscopy-derived protein ensembles deposited between January 1 and December 31, 2022 (Supplementary B.1.1). As a baseline, we compare against the landmark biomolecular modeling method AlphaFold3 (AF3) [15]. OC2 achieves a comparable precision score to AF3 in terms of root mean square deviation (RMSD) (Fig. 2a, left), confirming its advanced structure prediction accuracy. Regarding recalling all known ensemble states, OC2 outperforms AF3 (See Supplementary Fig. C1a for OC2’s sampling results). This advantage stems from the fundamental difference between the two models: AF3 is optimized for static structure prediction while OC2 models biomolecules as a conformational landscape to capture the underlying dynamic behaviors. Notably, OC2 also continues to improve in RMSD recall as sampling size increases, whereas AF3 plateaus due to lower prediction diversity (Fig. 2b, right). The precision scores of both methods remain stable with increased sampling, indicating that greater conformational diversity does not compromise structural accuracy. OC2 quantitatively recover protein flexibility, as measured by the average root mean square fluctuation (RMSF) across all backbone atoms, which closely matches NMR’s result (1.20 Å vs. 1.51 Å). In contrast, AF3 predictions exhibit lower flexibility (0.54 Å RMSF). OC2 also achieves higher correlation with experimental RMSF values, outperforming AF3 in both per-target and global Pearson correlation coefficients (Fig. 2a, right). Similar trends are observed when assessing distributional accuracy using cosine similarity between the top principal components of predicted and true ensembles, further validating OC2’s excellent ability to generate thermodynamic ensembles.

**Fig. 2.**
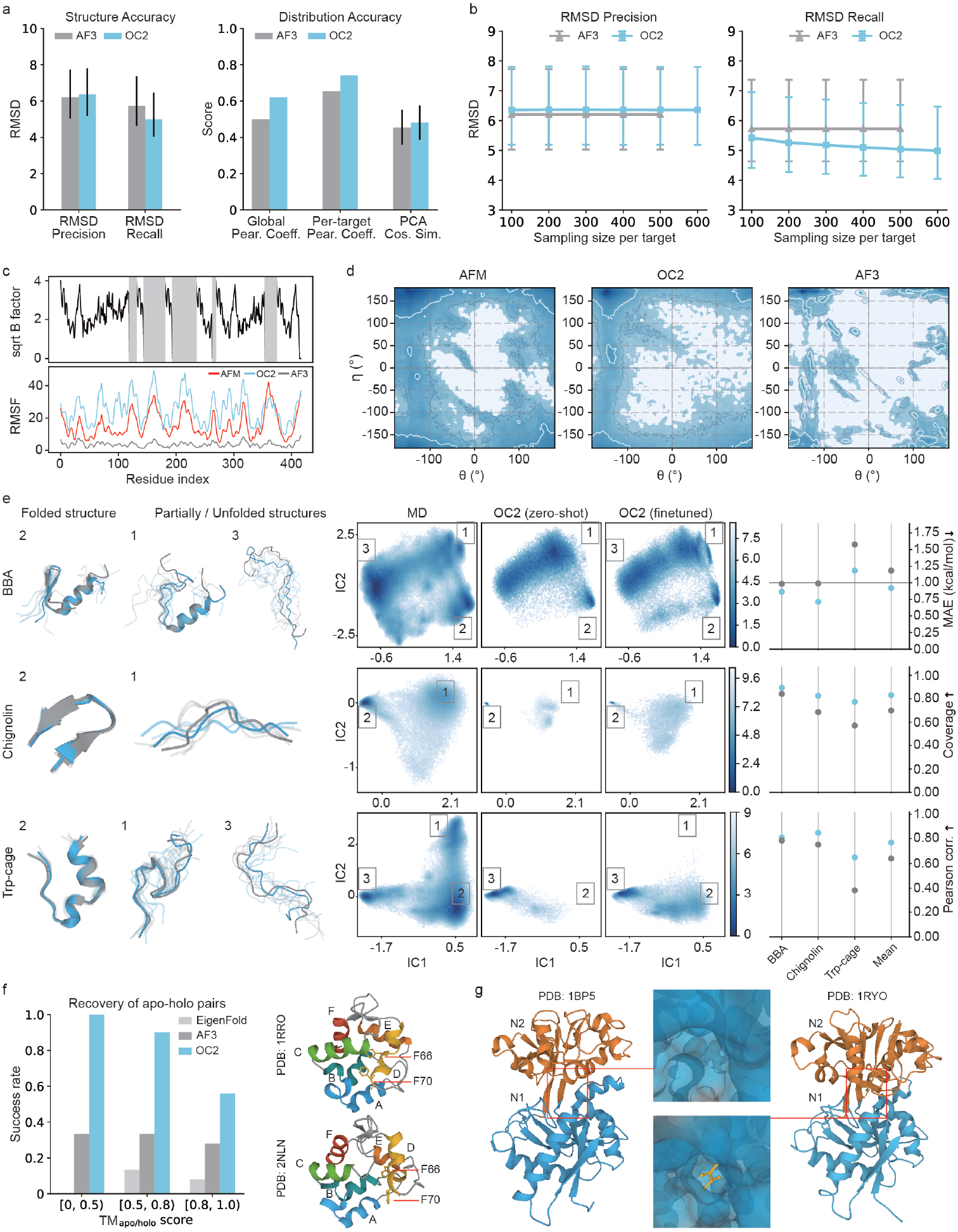
OC2 effectively samples biomolecular conformational distributions. Error bars indicate 95% confidence intervals. **a**, Performance comparison on 50 NMR ensembles using structural (left) and distributional (right) metrics. Precision: the average RMSD from each predicted structure to its closest crystal structure. Recall: the average RMSD from each crystal structure to its closest predicted structure. **b**, RMSD precision and recall scores on the same NMR ensembles as a function of sampling size. **c**, Distribution recovery of RNase P (PDB ID: 2A64). Bottom: Per-residue RMSF of RNase P conformers sampled from AFM, AF3, and OC2. Top: B-factors from the RNase P PDB structure serve as a reference (square root form applied to maintain physical consistency with RMSF). **d**, Conformational distributions of AFM, AF3, and OC2 in pseudo-torsion angle space. **e**, Comparison of fast-folding protein distributions sampled by the DESRES Anton supercomputer vs. OC2 models either trained on PDB structures or fine-tuned on DESRES fast folders. Left: Folded, partially folded, and unfolded structures predicted by OC2 (light blue) compared to MD ground truth (gray). Middle: Free energy landscapes (kcal/mol) from MD and OC2 in the space of the two slowest TICA dimensions. Right: comparison of free energy landscapes in terms of MAE of free energy differences (first row), coverage of the landscape (second row) and Pearson correlation coefficients of free energy. **f**, Performance in recovering alternative states from an apo/holo benchmark of 83 protein pairs. Left: Recovery success rate across pairs binned by pairwise structural similarity (**TM**_apo*/*holo_). Right: Example of a Ca^2+^-induced apo/holo transition. Helices are labeled and colored in different colors. **g**, Ligand-induced cryptic pocket formation in the N-lobe of Human Serum Transferrin.

Next, we demonstrate that OC2 can accurately reproduce RNA conformational ensembles, where structural flexibility is crucial for substrate specificity and functional regulation. As a stringent test case, we consider Ribonuclease P (RNase P, PDB ID: `2A64`), a ubiquitous trans-acting RNA enzyme that processes the 5’ ends of precursor tRNA [25]. Lee et al. [26] recently recovered the full conformational landscape of fulllength RNase P using atomic force microscopy (AFM) and machine learning, releasing an ensemble of 1,578 representative structures. These structures exhibit wild conformational fluctuations (with RMSF amplitudes up to 60 Å for certain coordinates), presenting a significant challenge for predictive models. To benchmark performance, we generate matching ensembles of 1,578 structures with both OC2 and AF3. OC2 closely reproduces the experimental per-base RMSF profile, achieving a Pearson correlation of 0.77 with the AFM reference (Fig. 2c). In contrast, AF3 predictions collapse onto a single dominant conformation, displaying minimal variability. We further compare pseudo-torsion-angle distributions between both methods. As shown in Fig. 2d, OC2 spans a significantly broader range that better matches experimental observations compared to AF3. These results underscore OC2’s potential in revealing conformational heterogeneity of RNAs

#### 2.1.2 Matching MD equilibrium distributions

As a quantitative investigation, we evaluate OC2’s ability to reproduce free energy landscapes obtained from extensive MD simulations. Accurately characterizing biomolecular conformations and estimating free energy landscapes via MD typically requires hundreds of microseconds to milliseconds of simulation time, making such simulations feasible only in limited cases due to immense computational costs [7]. For benchmarking, we leverage millisecond-scale D. E. Shaw Research (DESRES) simulations of 12 fast-folding proteins [7] generated on the Anton supercomputer, following the evaluation framework of Lewis et al. [27]. Most training structures in the PDB correspond to low-energy states, providing limited supervision for transition states in the free energy landscape. To mitigate this issue, we fine-tune a variant of OC2 on 9 out of the 12 fast-folding proteins, reserving the remaining 3 targets for evaluation (see Supplementary B.1.2 for details). For evaluation, we project the sampled conformations into a two-dimensional manifold using time-lagged independent analysis (TICA) [28].

Despite the absence of training signals for fine-grained conformational landscapes, the original OC2 model generates meaningful conformational distributions and identifies critical metastable states (Fig. 2e, middle). Quantitatively, the average free energy mean absolute error (MAE) between the MD and model 2D free energy landscapes in the TICA space is only 1.18 kcal/mol (Fig. 2e, right), on the order of difference between two classical force fields [29]. Additionally, OC2 samples effectively cover around 70% of conformations sampled by simulations and exhibit an average Pearson correlation of 0.65 for free energy across the TICA space (Fig. 2e, right). Fine-tuning on nine proteins yields notable improvement, enabling OC2 to more accurately recover the free energy landscape, especially for those low-density regions. Across all tested proteins, the fine-tuned OC2 model accurately predicts the native, unfolded, and intermediate states (Fig. 2e, left). Quantitatively, it attains an average free energy MAE of 0.92 kcal/mol and improves the coverage and Pearson correlation to 83% and 0.77, respectively (Fig. 2e, right).

OC2 also demonstrates close agreement with MD simulations in predicting secondary structure propensities (Supplementary Fig. C1b, right). Even without direct fine-tuning on MD data, OC2 achieves a significantly closer alignment with MD-derived equilibrium distributions than AF3 (Supplementary Fig. C1), further underscoring its advantage to model conformational landscapes.

#### 2.1.3 Capturing conformational changes related to protein function

We further evaluate OC2 in terms of capturing critical conformational changes reflecting therapeutically relevant functions. We first examine induced-fit transitions, where binding interactions drive significant conformational changes in proteins. These transitions promote key regulatory mechanisms, including signal transduction, cooperativity, and allosterism. For benchmarking, we adopt a curated benchmark of apo/holo pairs associated with ligand-induced conformational changes [30], where most targets are released before October 1, 2021. OC2 significantly outperforms baseline methods—including AF3 and EigenFold [31]—in recovering both apo and holo states (Fig. 2f, left). Notably, the performance gap is particularly pronounced for proteins with highly distinct functional states (Supplementary Fig. C1c presents a detailed comparison in terms of TM-score).

Illustratively, we use OC2 to investigate the structural transitions of rat *β*- parvalbumin, a protein that undergoes substantial structural rearrangements upon binding (or release) of Ca^2+^. As shown in the right column of Fig. 2f, OC2 accurately captures both Ca^2+^-bound and Ca^2+^-free conformations. Specifically, in the Ca^2+^-free state, helix *D* undergoes significant displacement, reducing its contact with helices *A* and *B*. According to Henzl and Tanner [32], in the Ca^2+^-bound protein *F* 70 is intimately associated with apolar side chains from the *AB* domain, anchoring the C terminus of helix *D* to the *AB* domain. Our results also show that *F* 70 is deeply buried in the protein interior in the Ca^2+^-bound state. Whereas in the Ca^2+^-free state, *F* 70 has withdrawn from the hydrophobic core, indicating a loss of interdomain association mediated by the Ca^2+^. A similar displacement is observed for *F* 66.

Next, we consider cryptic pockets, which often emerge during protein dynamics. Understanding the formation of these transient binding sites is crucial for drug discovery, as it can reveal druggable pockets not present in static structures, potentially transforming previously ‘undruggable’ proteins into viable therapeutic targets. As a well-known case of cryptic pocket, Human Serum Transferrin functions through a dynamic process involving binding, transporting and releasing ions in specific sites [33]. To showcase OC2 ‘s sampling ability, we analyze the N-lobe (amino-termina) of Human Serum Transferrin that comprises two subdomains connected by a hinge. With the close motion of the hinge, a deep cleft is generated to contain the iron binding site [33, 34]. OC2 successfully captures this intricate conformational transition, effectively modeling the cryptic pocket’s formation and interruption (Fig. 2g, right).

### 2.2 Structure Prediction Performance

Although OC2 is primarily designed to characterize conformational distributions, it can be naturally extended to predict static structures by identifying the most representative conformation. Here, we evaluate OC2 ‘s structural accuracy on several standard prediction benchmarks. For evaluation, we extract representative conformations using an averaging-based approach (Supplementary B.2).

#### 2.2.1 Predicting Biomolecular Structures and Interactions

Considering OC2 ‘s distinct advantage in modeling conformational variability, we first evaluate its performance on highly flexible single-chain biomolecular structures, such as RNAs. In Fig. 3a, OC2 outperforms AF3 on challenging RNA targets presented in CASP15. Supplementary Fig. C2a,d,e provides additional analysis, showcasing specific CASP15-RNA examples where OC2 delivers more accurate predictions. Further analysis of RNA-only target prediction accuracy is presented in Supplementary Fig. C2c, where OC2 also exhibits a performance advantage. As the number of sampled structures increases, OC2 progressively captures higher-quality conformations (Supplementary Fig. C2b). We also evaluate OC2 on protein monomer datasets, where it achieves accuracy comparable to AF3 (Fig. 3b).

**Fig. 3.**
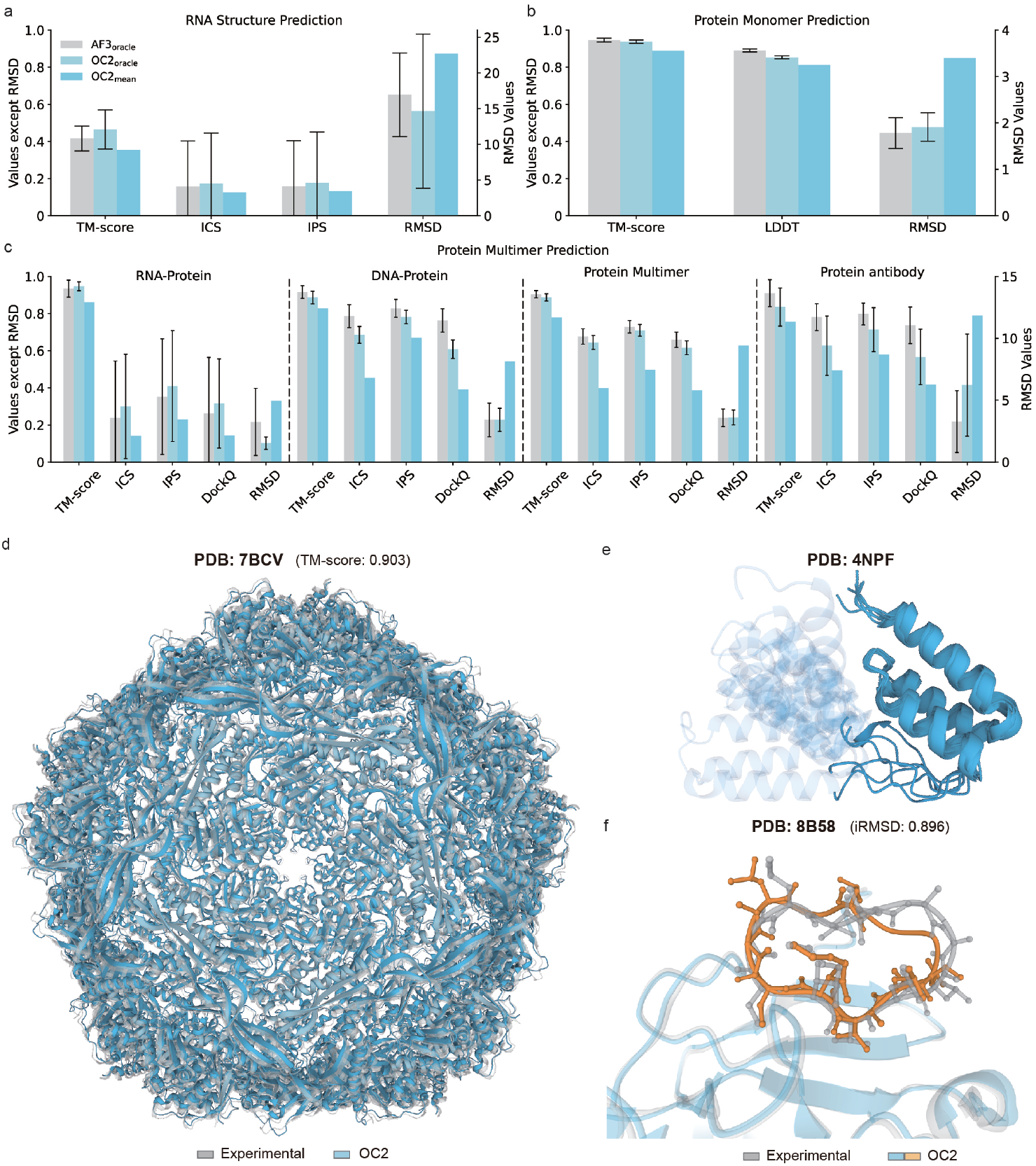
OC2 accurately predicts biomolecular structures and interactions. Error bars indicate 95% confidence intervals. **a**-**c**, Performance on diverse biomolecular datasets. All metrics represent the oracle value calculated from 25 samples using the OpenStructure Suite [35], with details provided in Supplementary B.2.3 **a**, Performance on CASP15 RNA targets, OC2 demonstrates higher accuracy than AF3. **b**, For single-chain proteins, OC2 matches the accuracy of AF3. **c**, In multi-chain biomolecular complexes, OC2 remains competitive performance in global metrics (*e*.*g*., RMSD). **d**,**e**,**f**, Biomolecular structures predicted by OC2. **d**, Icosahedral encapsulin from Brevibacterium linens, capable of protein compartmentalization for enzymatic reactions and iron storage (PDB ID: `7BCV`, 15,960 residues; TM-score: 0.903). **e**, Example of OC2 demonstrates flexibility in the loop region, a linker that connects two domains in the structure (PDB ID: 4NPF). **f**, The cyclic peptide Cyclosporin A contains several uncommon residues and is bound to Cyclophilin TgCyp23 from Toxoplasma gondii (PDB ID: `8B58`; interface RMSD (iRMSD) = 0.896 Å).

We further evaluate OC2 on a range of multi-chain complexes (Fig. 3c). OC2 outperforms AF3 on RNA–protein complexes, achieving lower RMSD (2.91 Å vs. 5.68 Å) and higher TM-score (0.94 vs. 0.85). For DNA–protein complexes, OC2 also shows better global structure metrics than AF3 (RMSD: 4.05 Å vs. 4.90 Å). On protein–protein complexes, both methods perform robustly, with AF3 showing slightly better results in interface metrics like DockQ (0.61 vs. 0.58). Currently, OC2 performs less effectively than AF3 on antibody structure prediction (DockQ: 0.57 vs. 0.68). These results highlight the complementary strengths of the two approaches: OC2 shows advantages on flexible systems such as nucleic-protein complexes, while AF3 excels at modeling highly ordered interfaces.

#### 2.2.2 Scaling to Model Large-scale Assemblies

Accurately modeling ultra-large biomolecular assemblies remains a major challenge due to the exponential increase in computational complexity with system size. OC2 addresses this by incorporating symmetry-aware optimizations, enabling efficient and accurate large-scale structure prediction. We demonstrate that OC2 effectively scales to predict large symmetric assemblies, including those with icosahedral (I), octahedral (O) and tetrahedral (T) symmetry. Fig. 3d highlights successful prediction of an icosahedral encapsulin from Brevibacterium linens (PDB ID: `7BCV`), a protein nanocompartment known for its role in enzymatic reactions and iron storage. This structure comprises 60 homologous chains, totaling 15,960 amino acids and forming 150 protein-protein interfaces. Quality assessment indicates consistently accurate interface predictions, with an average DockQ score of 0.577 and interface RMSD values 2.223 Å. Accurate predictions are also achieved for assemblies with O and T symmetry (Supplementary Fig. C2g). Additionally, high-quality viral protein models with TM-scores above 0.9 are presented in Supplementary Fig. C2f, highlighting OC2 ‘s strength in modeling large macromolecular structures.

### 2.3 Small Molecule Prediction

Building on our protein structure distribution modeling approach, we next explore how OC2 applies to small molecule prediction. In computational drug discovery, the prevalent focus on predicting a single “correct” ligand pose can overlook that multiple binding modes often exist with comparable probabilities. This is particularly relevant in early-stage discovery where protein-ligand interactions are not yet optimized.

#### 2.3.1 Capturing the ligand docking site with high validity

We first evaluate OC2 ‘s ability to predict protein-ligand interfaces on a subset of the PoseBusters Version 2 (PB-V2) benchmark comprising 213 targets without data leakage (Supplementary B.3.1). Following standard protocols, we measure accuracy as the fraction of predictions with pocket-aligned ligand RMSD to ground truth below 2 Å [36]. With 25 samples, OC2 achieves a 77% success rate given only protein sequence and ligand chemical composition (Fig. 4a).

**Fig. 4.**
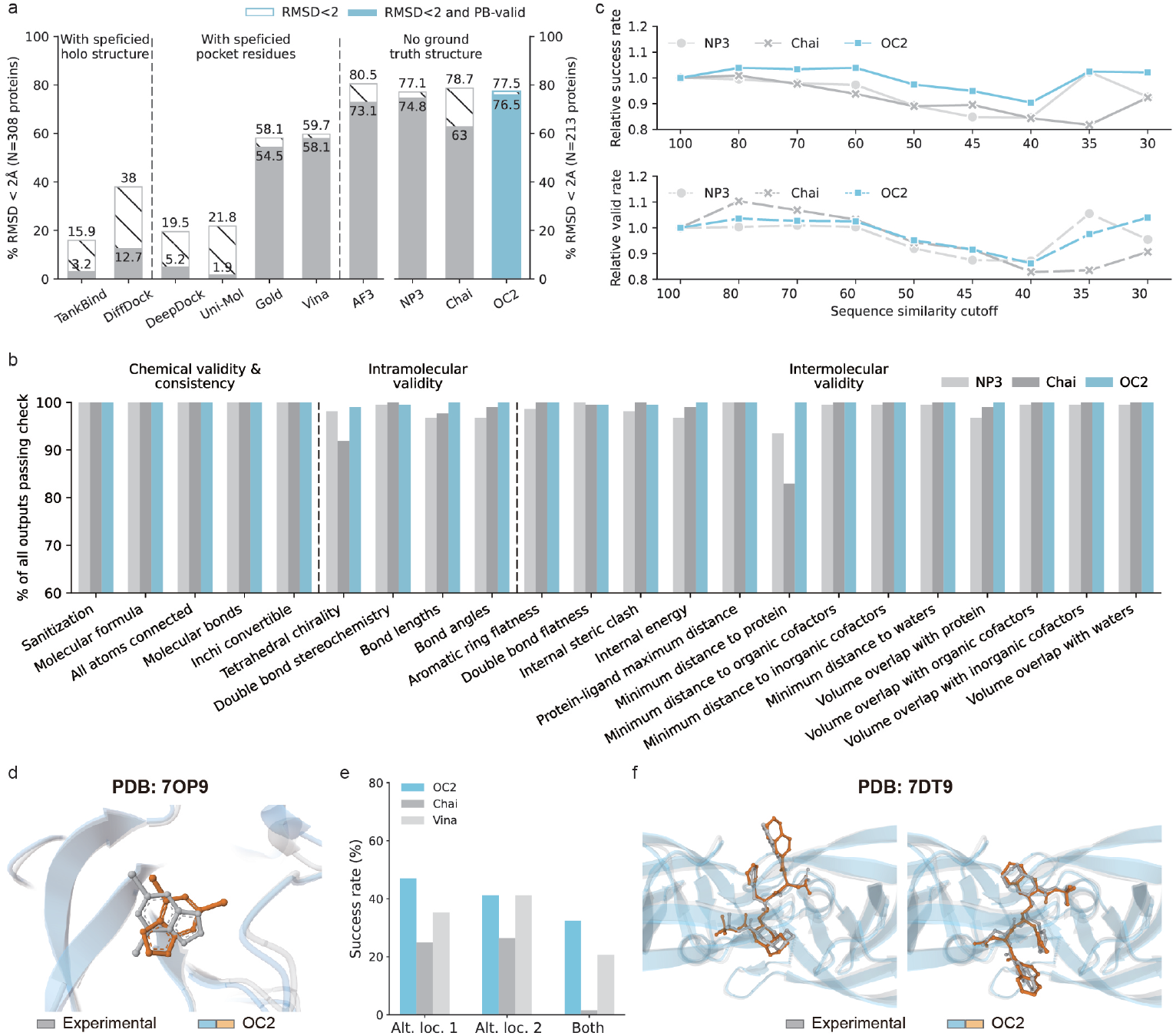
Ligand accuracy and validity analysis. **a**, Comparison of OC2 and baseline methods on the PB-V2 benchmark set with 213 protein-ligand structures. The value of OC2 is reported from the oracle value calculated from 25 samples, and the values of other tools except Chai and NP3 are collected from the PoseBusters prediction results published by AF3. **b**, Detailed validity check comparison on 213 PB-V2 targets. **c**, Relative score of success and valid rates across different protein sequence similarity cutoffs. **d**, OC2 ‘s predicted ligand is situated within the correct pocket and has passed the PB-valid check, yet the RMSD exceeds 2 Å due to a molecular rotation phenomenon. **e**, Success rate of OC2, Chai, and Vina in predicting ligands with multiple alternative locations. **f**, OC2 successfully predicts two different occupancies of ligand in PDB entry `7DT9`.

Beyond positional accuracy, the stereochemical validity of predicted ligands is crucial for real-world applications. We applied the PoseBusters quality-check pipeline (Supplementary B.3.1) and found that OC2’s predictions exceed those of NP3 and Chai in overall validity, though OC2 shows slightly weaker performance in specific metrics like double bond stereochemistry (Fig. 4b). Notably, when examining performance across different protein sequence similarity thresholds between PB-V2 and pre-2019 PDB proteins, OC2’s accuracy increases as sequence similarity decreases, while competitors show the opposite trend (Fig. 4c), demonstrating superior generalization to novel sequences. We found that in some cases, the ligands predicted by OC2 were located in the correct pockets and passed the PB validity test, yet the RMSD exceeded 2 Å due to the molecular rotation phenomenon (Fig. 4d). To gain a more comprehensive understanding, future investigations should incorporate advanced methodologies, such as free energy perturbation (FEP) calculations, to systematically compare the energetic differences among these conformations.

#### 2.3.2 Identifying Ligands with Multiple Docking Sites

Consistent with our distribution modeling approach, OC2 naturally extends to sampling multiple potential binding modes. We tested this on 68 small molecule targets with experimentally verified multiple binding sites (Supplementary B.3.2). By calculating pocket-aligned ligand RMSD, OC2 reaches 47.1% prediction accuracy for alternative locations 1 and 41.2% for alternative locations 2 at the cutoff of 2 Å, significantly outperforming Chai with 25% and 26.5%, respectively. OC2 successfully predicts both locations for 22 targets, while Vina Dock and Chai only predict 14 and 1, respectively (Fig. 4e).

This capability emerges naturally from OC2’s structural distribution modeling. For example, with PDB entries shown in Fig. 4f, OC2 successfully identifies multiple experimentally observed binding locations, and the distance between the two positions of the experimental ligand does not correlate with OC2’s predictive ability (Supplementary Fig. C2j). This ability to explore broader conformational landscapes aligns with our overall framework’s design principles and offers practical advantages for drug discovery, where considering multiple binding configurations can enhance lead optimization.

## 3 Current Limitations & Future Directions

OC2 marks a substantial step toward statistically understanding biomolecular conformational landscapes, yet several limitations remain and motivate our ongoing research.

### Scope of Applicability

Although OC2 demonstrates generalizability across the protein and RNA test cases presented in this report, its performance on more heterogeneous systems is still uncharted. Examples include RNA–DNA hybrids, multicomponent nucleoprotein assemblies, and very large macromolecular machines characterized by extensive conformational variability. Rigorous experimental benchmarks for these biomolecules are required to specify the current application boundary of OC2. Therefore, establishing such benchmarks and extending OC2 on these systems serves as the top priority in our future work.

### Emulating equilibrium distributions of more complicated systems

Although OC2 attains near–chemical accuracy on fast-folding test systems (evidenced by the average free energy MAE of 0.92 kcal/mol in Fig. 2) and faithfully reproduces NMR-derived ensembles, its thermodynamic fidelity for more demanding cases remains unproven. Large multi-domain proteins, assemblies with multiple interaction partners, and systems whose energetics depend on experimental variables such as temperature or pH will require dedicated validation. Another open challenge is the efficient sampling of rare but functionally essential substates that are separated by high free-energy barriers.

### Understanding divergence from SOTA static structure predictors

Benchmarking shows that OC2 can yield structures that diverge from SOTA static structure predictors such as AF3, especially for antibody–antigen complexes. This discrepancy possibly arises from OC2’s ensemble-centric philosophy: it prioritizes capturing conformational heterogeneity rather than converging on a single “best” structure. While this can reveal alternative binding modes and mechanistic insights, it may also produce conformations that deviate from crystallographic interfaces in PDB. Resolving these differences will require objective evaluation protocols and close collaboration with wet-lab experiments.

### Integrate OC2 into real-world applications

OC2 already exhibits capability in generating low-energy conformers and tracking conformational transitions of druglike molecules in protein–ligand environments, but leveraging these strengths for realworld drug discovery demands further work. Systematic benchmarks are necessary to assess how accurately OC2 reproduces experimental conformational energy differences, entropic contributions, and low-population states that drive pharmacological activity. Comparative studies against rigorous baselines—FEP, advanced enhanced-sampling protocols, and quantum-chemical calculations—are essential to position OC2 within existing workflows.

## Appendix A Training Details

### A.1 Training Dataset

We constructed a generative foundation model for learning biomolecular conformation distributions, primarily trained on data from the RCSB Protein Data Bank (PDB)—a comprehensive repository of experimentally determined protein structures. For structure distribution modeling, we curated a broad dataset comprising all experimental structures available in the PDB prior to September 30, 2021, applying a date filter to prevent data leakage while deliberately retaining entries of all resolutions and experimental methods to capture the full diversity of structural data.

To ensure reliable model evaluation, we applied filtering criteria to curate highquality training targets for biomolecular structure prediction tasks. Specifically, we excluded NMR entries and selected only crystal structures with resolution better than 9 Å to maintain structural quality and consistency. To reduce bias between training and evaluation sets, we performed clustering on PDB chains and interfaces. Chainlevel clustering was conducted at 40% sequence homology for proteins, 100% homology for nucleic acids (only identical sequences were grouped into the same cluster), and 100% homology for peptides (fewer than 10 residues). Interface-level clustering was performed by joining the cluster IDs of constituent chains. Interfaces *I* and *J* were assigned to the same interface cluster *C*_interface_ only if their constituent chain pairs *{I*_1_, *I*_2_*}, {J*_1_, *J*_2_*}* shared the same chain cluster pairs *{C*_chain1_, *C*_chain2_*}*.

#### A.2 Data Pipeline

The initial structural features provided to the OC2 data pipeline offer a comprehensive representation of molecular topology, including atom, residue and motif level features. For ligands, non-standard residues, and post-translational modifications (PTMs), all atomic details are preserved to maintain structural integrity and capture key interactions critical for downstream analysis.

OC2 leverages MSA-based conditioning to enhance structure prediction capabilities. During model training, MSAs are generated using jackhmmer [37] against databases UniRef90 [38], MGnify [39], and reduced version of the BFD [13, 40]; hhblits [41] searches the full BFD; and nhmmer [42] retrieves MSAs from Rfam [43], RNACentral [44], and the Nucleotide collection [45]. Additionally, MMseqs [46] clusters both protein and RNA sequence databases. No templates are used in our pipeline to ensure flexibility in processing diverse inputs, allowing the model to generalize effectively across various scenarios without being constrained by predefined patterns.

## Appendix B Evaluation Methods

### B.1 Protein Distribution Prediction

Here, we detail our evaluation framework for assessing OC2’s performance in predicting conformational distributions.

#### B.1.1 Reproducing thermodynamic ensembles

As described in Sec. 2.1.1, we evaluate OC2’s ability to reproduce thermodynamic protein ensembles that were not included in the training data. Specifically, we construct an evaluation benchmark using 50 NMR ensembles deposited in the PDB between January 1, 2022, and December 31, 2022. For the results in Fig. 2a-b, we generate 500 conformers per protein using OC2 and AF3. To assess how well the predicted ensembles recover known NMR structures, we compute precision and recall scores using RMSD. To assess distributional accuracy, we calculate residue-level RMSF across the sampled ensembles and measure the Pearson correlation between OC2/AF3 predictions and NMR ground truth—both globally (across all proteins) and per target. Following Jing et al. [22], we also compute the average unsigned *cosine* similarity between the top principal components of the predicted and experimental ensembles.

For the RNA evaluation in Fig. 2c-d, we sample 1,578 conformers with OC2 and AF3 for RNase P and compare per-base RMSF and pseudo-torsion angle distributions against the 1,578 AFM-resolved conformational states [26]. Specifically, we compute RMSF using the C4’ atom of each base and visualize pseudo-torsion angle distributions using 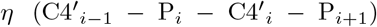 and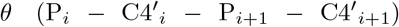, where *i* indexes RNA bases. The AFM-resolved conformational data are obtained from the official repository (https://home.ccr.cancer.gov/csb/pnai/data/conformationalspace/ConfspaceRNasePRNA/DNN/Top10models).

##### B.1.2 Matching MD equilibrium distributions

As described in Sec. 2.1.2, we test OC2’s ability to recover equilibrium distributions derived from extensive MD simulations. Our benchmark consists of 12 fast-folding protein systems simulated with the CHARMM22* force field, totaling 8.2 *ms* of cumulative simulation time [7]. Starting from the OC2 model pretrained on general biomolecular structures in PDB, we then fine-tune the model on 9 of 12 fast-folding proteins and test the performance of both the pretrained (denoted as ‘OC2 (zeroshot)’) and fine-tuned (denoted as ‘OC2 (finetuned)’) models on the remaining 3 proteins. The training set includes BBL, Homeodomain, NTL9, Protein B, Protein G, Villin, WW, a3D, and Repressor, while the test set consists of BBA, Chignolin, and Trp-cage.

For the results presented in Fig. 2e, we sample 100,000 conformations per protein. Following Lewis et al. [27], we assess the similarity between OC2-sampled distributions and MD simulations by comparing representative structures, free energy landscapes, and secondary structure propensities. To facilitate this comparison, we apply timelagged independent component analysis (TICA) [28] to project the conformations onto a low-dimensional manifold, using pairwise C_*α*_ distances as features. The standard TICA projection is performed with PyEmma [47], using a lag time of 20 *ns*. Representative structures from MD simulations are obtained via clustering analysis, with the first 16 TICA components serving as clustering features. The free energy landscape is then visualized in the space of the two slowest TICA dimensions. For evaluating the generated free energy landscapes in the space of the two slowest TICA components, we evenly split the landscape into 50 *×* 50 bins. On those bins with MD density larger than a predefined threshold, we calculate the MAE and the Pearson correlation coefficients of the free energy differences between MD and OC2, as well as the coverage ratio of OC2’s conformations over MD reference. The implementation is inspired by the evaluation codebase of BioEmu(https://github.com/microsoft/bioemu-benchmarks) [27]. For secondary structure propensity calculations shown in Supplementary Fig. C1b, we use the experimentally determined structures or ensembles from the corresponding PDB entries as ground truth (denoted as ‘PDB’). Additionally, we sample 50,000 conformers for the same evaluation proteins using AF3 (‘AF3 (zero-shot)’) and compare its predictions with those of the pretrained OC2 model. The results are summarized in Supplementary Fig. C1b. Notably, we observe that increasing the number of AF3 samples does not improve its performance, so we limit AF3 sampling to 50,000 conformers to conserve computational resources.

##### B.1.3 Capturing conformational changes to protein function

To evaluate OC2’s ability to sample functionally relevant conformational changes, we examine two biologically significant benchmarks: a dataset of 88 apo/holo protein pairs associated with ligand-induced conformational changes from Saldaño et al. [30], and a cryptic pocket benchmark comprising 13 protein pairs [27].

Following Jing et al. [31], we assess OC2’s performance in capturing multiple conformational states by computing TM-score between model predictions and corresponding ground-truth structures, using an ensemble TM-score to quantify accuracy of predicting diverse states:

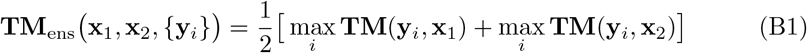

where **x**_1_, **x**_2_ are ground truth structures and *{***y**_*i*_*}* are OC2 samples. A naive baseline model always predicting a single conformation serves as reference, denoted as **TM**_apo*/*holo_. During inference, we include all available biomolecular partners to consider all potential environmental effects. Both the AF3 web server and local program accept limited valid ligands as input, so we skip the ligand after an attempt for such cases. For results in Fig. 2f (left), we report success rate as the fraction of cases where OC2 outperforms this baseline. A detailed TM-score comparison is provided in Supplementary Fig. C1b. For both OC2 and AF3, we generate predictions using the structural assembly corresponding to the first PDB entry, following the convention established by AF3 [15].

#### B.2 Biomolecular Structure Prediction

We evaluated OC2’s performance in biomolecular structure prediction using several benchmark datasets including proteins, nucleic acids, biomolecular interactions, and ultra-large proteins, employing well-established metrics for assessment.

##### B.2.1 Predicting Biomolecular Structure and Interactions

We conducted a comprehensive evaluation using a carefully curated dataset to assess the model’s ability to generalize to novel structures. This dataset derives from 9,612 PDB structures released between May 30, 2022 and January 12, 2023—substantially after the training cutoff dates of both OC2 and AF3. To ensure minimal homology to training data, we applied a rigorous two-stage clustering process. We clustered protein sequences in the PDB based on sequence identity using mmseqs easy-cluster. The structures in our test dataset combined with our training set (182,421 structures) were clustered at 40% sequence identity, producing 2,598 structures with distinct cluster IDs. We then selected protein sequences from chains not part of training set clusters, excluding proteins with resolution greater than 4.5 Å to maintain high structural quality.

To better investigate the accuracy for nucleic acids, ten RNA targets publicly available on CASP15 website were selected for evaluation to ensure a comprehensive assessment of the model. It includes `R1107/7QR4`, `R1108/7QR3,R1116/8S95`, `R1117/8FZA`, `R1128/8BTZ`, `R1136/7ZJ4`, `R1149/8UYS`, `R1156/8UYE`, `R1169/7YR7`, `R1170/7YR6`. Due to the limited dataset size of CASP15-RNA, we further assess the precision of predicting nucleic acids alone, independent of protein interactions. The PDB ID list is as follows: `7MKT`, `7PNL`, `7QVQ`, `7R6N`, `7TZT`, `7TZU`, `7UCR`, `7URI`, `7URM`, `7VFT`, `8D28`, `8D2A`, `8D2B`, `8D5L`, `8D5O`.

##### B.2.2 Scaling to Model Large-scale Assemblies

Current structure prediction methods like AF3 struggle with structures exceeding 5,120 residues. We selected representative cases—including those with icosahedral symmetries—to evaluate our method’s capability in handling extremely large macromolecular systems. One example is an encapsulin from Brevibacterium linens that enables protein compartmentalization for enzymatic reactions and iron storage, consisting of 60 homologous chains totaling 15,960 amino acids and forming 150 interfaces (PDB ID: `7BCV`). Since many viruses adopt icosahedral (I) symmetry, we also assessed OC2 on representative viral capsids including `7TJE`, `7TJD`, `7TJG`, `8XEG`, `8TU1`, `8TU0`, `8TEX`, `7U94`, and `8FQ4`. Additionally, we evaluated structures with octahedral (O) and tetrahedral (T) symmetry, such as `7Q5Q`, `7UOL`, `7RVB`, `8IC8`, `8PVE`, and `7RAB`.

##### B.2.3 Structure Prediction Metrics

Following protocols similar to CASP for fair method comparison, we generated 25 samples for each method and evaluated the best-performing (oracle) structure across all metrics. Since OC2 models ensemble behavior of biomolecular structures, predicted samples naturally exhibit variability, especially in segments without stable secondary structure. This variability can reduce scores in metrics sensitive to local distance errors, such as LDDT.

To address this, we employed an averaging strategy to reduce sample variance and approximate the maximum likelihood structure. During reverse diffusion, we collected multiple graph representations along the trajectory, computed their average over a subset of samples, and fed this averaged representation into the structure solver to obtain a representative structure with improved metric scores.

We evaluated biomolecular structure prediction accuracy using OpenStructure Suite [35] (version 2.9.1) with metrics categorized into biomolecular-level and interfacelevel metrics. Biomolecular-level metrics such as Root Mean Square Deviation (RMSD) [48], Local Distance Difference Test (LDDT) [49], and Template Modeling score (TM-score) [50] were computed over the entire biomolecular structures. Interface-level metrics such as Interface Contact Similarity(ICS) [51], Interface Patch Similarity(IPS) [51], and DockQ [52] were evaluated specifically at protein–protein and protein–nucleic acid interfaces. Here, we define a complex’s interface metric as the mean of its individual interfaces.

RMSD quantifies the average atomic displacement between predicted and reference structures after optimal superposition, with lower values indicating higher similarity, but may be sensitive to outlier regions. LDDT assesses local structural accuracy by comparing nearby atomic distances without requiring superposition. TM-score measures global topological similarity, emphasizing larger-scale structural agreement. ICS evaluates the similarity of residue-residue contacts at the binding interface, while IPS assesses the geometric overlap and spatial consistency of interface patches. Finally, DockQ provides a score between 0 and 1, reflecting the overall quality of predicted protein-protein complexes through integrated structural and interface evaluation.

#### B.3 Small Molecule Prediction

Here, we provide the details in evaluating OC2’s ability in predicting small molecules.

##### B.3.1 Ligand accuracy and validity

The PoseBusters Version 2(PB-v2) benchmark contains 307 available entries, with the obsolete entry `7D6O` ignored. To prevent data leakage, we retain only the 213 targets released after the training dataset cutoff of September 30, 2021. As baselines, we collect the NP3-predicted protein-ligand structures from NPBench Data (https://zenodo.org/records/14503936) and the Chai-predicted protein-ligand structures from the official-released technical report (https://chaiassets.com/chai1/paper/assets/posebusterspredictions.zip). As AF3 doesn’t provide predicted complex structures, we sourced its PB-v2 results from the paper [15].

We calculated pocket-aligned ligand RMSD using the NPBench repository (https://github.com/iambic-therapeutics/np-bench) and performed PoseBusters quality checks using the PoseBusters repository (https://github.com/maabuu/posebusters). Sequence similarity between the PB-V2 set and training set was measured with HHSearch, with relative scores of success and valid rates calculated at sequence identity cutoff of 100%.

##### B.3.2 Ligands with multiple docking sites

Data for ligands with multiple docking sites were screened from our test set using alternative location information. To prevent clashes between alternative locations, we kept only entries with ligand center of gravity distance greater than 2 Å.

For the Vina dock baseline, we employed experimentally resolved protein structures from the PDB as receptor models. Ligand conformations were generated using the RDKit [53] conformation generation algorithm, followed by conversion of both proteins and ligands to pdbqt format using AutoDockTools (ADT) [54]. The docking protocol used search boxes encompassing entire proteins with parameters `--num modes 1000` to generate 1000 samples, `--energy range 1000` to retain all top-ranked modes, and `--exhaustiveness 1000` to ensure thorough conformational sampling.

We also evaluated two additional ensemble docking strategies: (1) selecting the top 1000 conformations from 10 independent docking runs with varying initial ligand conformations, and (2) assembling an ensemble of 1000 conformations by collecting the top 100 ranked poses from each of 10 independent runs. Neither approach yielded significant improvements in docking accuracy compared to standard protocol.

## Appendix C Supplementary Figures

**Fig. C1.**
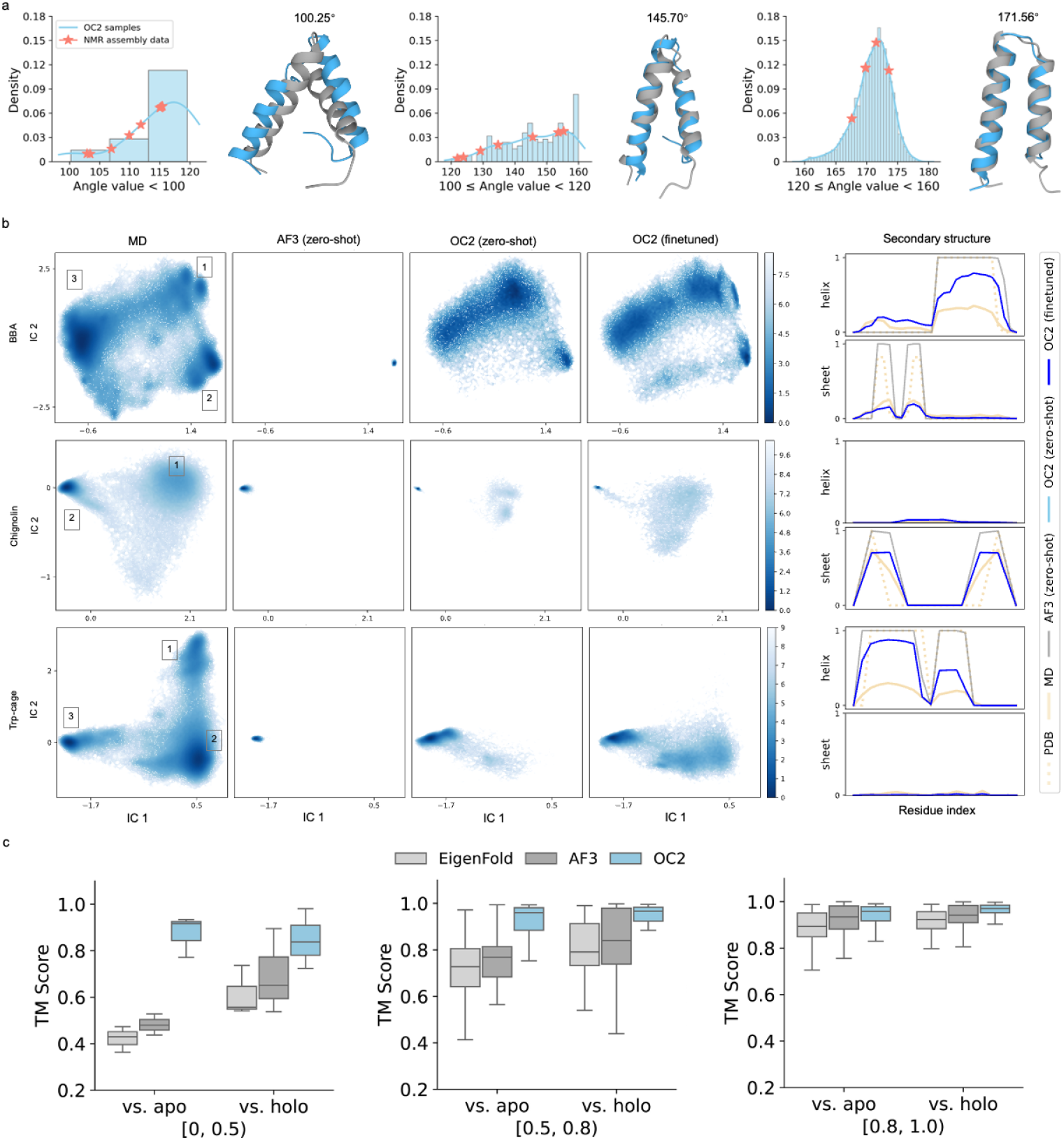
Additional results on distributional modeling. **a**, Case study on modeling the conformational distribution of NMR structure. We sample 6000 structures of a 45-residue NMR protein (PDB: 7LVG) containing two *α*-helices. The inter-helical angle distribution spans 100–180^*?*^, aligning well with the 20 NMR models (red asterisks), showing that the sampled conformations from OC2 broadly cover the experimental ensembles. Representative conformations at 100.25^*?*^, 145.70^*?*^, and 171.56^*?*^ match NMR structures with RMSDs of 5.92 Å, 3.63 Å, and 3.95 Å, respectively. **b**, Comparison of fast-folding protein conformational distributions sampled by OC2 and AF3, both trained on PDB structures. Left: Free energy surfaces (in kcal/mol) projected onto the two slowest TICA dimensions. Right: Secondary structure propensities comparison across the full structural ensemble. **c**, Performance on the apo/holo benchmark. Left to right: TM-score results across apo/holo subsets binned by structural similarity (**TM**_apo*/*holo_). OC2 shows a significant advantage over other methods, particularly when apo and holo states differ substantially.

**Fig. C2.**
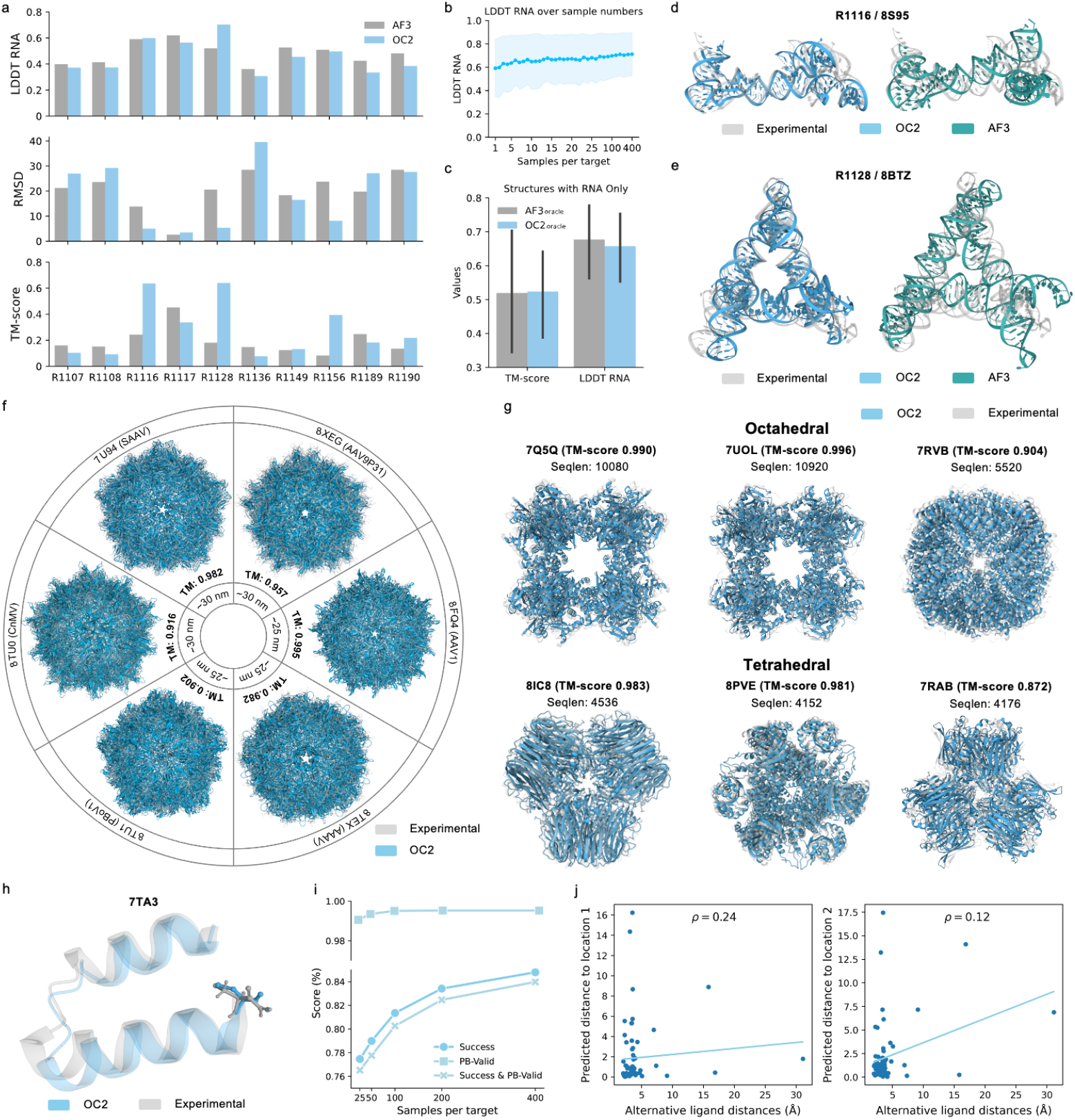
Additional results on biomolecular structure prediction. **a**. CASP15 RNA prediction accuracy from AF3 and OC2. The results presented here represent the oracle values derived from 25 individual samples taken from both AF3 and OC2. RNA LDDT was calculated with a 30 Å inclusion radius, rather than the standard 15 Å used for proteins, to accommodate their larger size. **b**. RNA prediction quality increases with the number of samples. The metrics (with 95% confidence intervals) of RNA structure prediction as a function of the number of samples. **c**. RNA-only prediction accuracy from AF3 and OC2. **d**. Case study of target R1116/8S95,the orange on the left represents AF3(RMSD:14.08 Å), the blue on the right represents OC2(RMSD:5.53 Å), and the gray represents native protein. **e**. Case study of target R1128/8BTZ, the orange on the left represents AF3(RMSD:15.12 Å), the blue on the right represents OC2(RMSD:6.52 Å), and the gray represents native protein. **f**. Representative examples of viral structure predictions, with TM-scores exceeding 0.9. **g**. Predicted structures with octahedral and tetrahedral symmetry. **h**. Accurate prediction of modified residues within polymer structures. **i**. The success rate increases with the sampling number per target, while the valid rate remains stable. **j**. The distance between the two ligand alternative positions correlates little with the predicted distance error.

## References

[1] Frauenfelder, H., Sligar, S.G., Wolynes, P.G.: The energy landscapes and motions of proteins. Science 254(5038), 1598–1603 (1991)

[2] Wlodawer, A., Vondrasek, J.: Inhibitors of hiv-1 protease: a major success of structure-assisted drug design. Annual review of biophysics and biomolecular structure 27(1), 249–284 (1998)

[3] Wei, G., Xi, W., Nussinov, R., Ma, B.: Protein ensembles: how does nature harness thermodynamic fluctuations for life? the diverse functional roles of conformational ensembles in the cell. Chemical reviews 116(11), 6516–6551 (2016)

[4] Shi, Y.: A glimpse of structural biology through x-ray crystallography. Cell 159(5), 995–1014 (2014)

[5] Danev, R., Yanagisawa, H., Kikkawa, M.: Cryo-electron microscopy methodology: current aspects and future directions. Trends in biochemical sciences 44(10), 837– 848 (2019)

[6] Marion, D.: An introduction to biological nmr spectroscopy. Molecular & Cellular Proteomics 12(11), 3006–3025 (2013)

[7] Lindorff-Larsen, K., Piana, S., Dror, R.O., Shaw, D.E.: How fast-folding proteins fold. Science 334(6055), 517–520 (2011)

[8] Shaw, D.E., Grossman, J., Bank, J.A., Batson, B., Butts, J.A., Chao, J.C., Den-eroff, M.M., Dror, R.O., Even, A., Fenton, C.H., et al.: Anton 2: raising the bar for performance and programmability in a special-purpose molecular dynamics supercomputer. In: SC’14: Proceedings of the International Conference for High Performance Computing, Networking, Storage and Analysis, pp. 41–53 (2014). IEEE

[9] Shaw, D.E., Adams, P.J., Azaria, A., Bank, J.A., Batson, B., Bell, A., Bergdorf, M., Bhatt, J., Butts, J.A., Correia, T., et al.: Anton 3: twenty microseconds of molecular dynamics simulation before lunch. In: Proceedings of the International Conference for High Performance Computing, Networking, Storage and Analysis, pp. 1–11 (2021)

[10] Laio, A., Parrinello, M.: Escaping free-energy minima. Proceedings of the national academy of sciences 99(20), 12562–12566 (2002)

[11] Yang, Y.I., Shao, Q., Zhang, J., Yang, L., Gao, Y.Q.: Enhanced sampling in molecular dynamics. The Journal of chemical physics 151(7) (2019)

[12] Joshi, S.Y., Deshmukh, S.A.: A review of advancements in coarse-grained molecular dynamics simulations. Molecular Simulation 47(10-11), 786–803 (2021)

[13] Jumper, J., Evans, R., Pritzel, A., Green, T., Figurnov, M., Ronneberger, O., Tunyasuvunakool, K., Bates, R., Žídek, A., Potapenko, A., et al.: Highly accurate protein structure prediction with alphafold. Nature 596(7873), 583–589 (2021)

[14] Baek, M., DiMaio, F., Anishchenko, I., Dauparas, J., Ovchinnikov, S., Lee, G.R., Wang, J., Cong, Q., Kinch, L.N., Schaeffer, R.D., Millán, C., Park, H., Adams, C., Glassman, C.R., DeGiovanni, A., Pereira, J.H., Rodrigues, A.V., Van Dijk, A.A., Ebrecht, A.C., Opperman, D.J., Sagmeister, T., Buhlheller, C., Pavkov-Keller, T., Rathinaswamy, M.K., Dalwadi, U., Yip, C.K., Burke, J.E., Garcia, K.C., Grishin, N.V., Adams, P.D., Read, R.J., Baker, D.: Accurate prediction of protein structures and interactions using a three-track neural network. Science 373(6557), 871–876 (2021) 10.1126/science.abj8754

[15] Abramson, J., Adler, J., Dunger, J., Evans, R., Green, T., Pritzel, A., Ronneberger, O., Willmore, L., Ballard, A.J., Bambrick, J., et al.: Accurate structure prediction of biomolecular interactions with AlphaFold3. Nature 630(8016), 493–500 (2024)

[16] ByteDance AML AI4Science Team, Chen, X., Zhang, Y., Lu, C., Ma, W., Guan, J., Gong, C., Yang, J., Zhang, H., Zhang, K., Wu, S., Zhou, K., Yang, Y., Liu, Z., Wang, L., Shi, B., Shi, S., Xiao, W.: Protenix - advancing structure prediction through a comprehensive AlphaFold3 reproduction. bioRxiv (2025) 10.1101/2025.01.08.631967

[17] Chai Discovery team, Boitreaud, J., Dent, J., McPartlon, M., Meier, J., Reis, V., Rogozhonikov, A., Wu, K.: Chai-1: Decoding the molecular interactions of life. bioRxiv (2024) 10.1101/2024.10.10.615955

[18] Wohlwend, J., Corso, G., Passaro, S., Reveiz, M., Leidal, K., Swiderski, W., Portnoi, T., Chinn, I., Silterra, J., Jaakkola, T., Barzilay, R.: Boltz-1 democratizing biomolecular interaction modeling. bioRxiv (2024) 10.1101/2024.11.19.624167

[19] Qiao, Z., Ding, F., Dresselhaus, T., Rosenfeld, M.A., Han, X., Howell, O., Iyengar, A., Opalenski, S., Christensen, A.S., Sirumalla, S.K., Manby, F.R., III, T.F.M., Welborn, M.: NeuralPLexer3: Accurate Biomolecular Complex Structure Prediction with Flow Models (2024). https://arxiv.org/abs/2412.10743

[20] Wayment-Steele, H.K., Filippova, O., Sergey, R., Rao, R.M.: Predicting multiple functional conformations for protein-protein complexes with a sequence-based model. bioRxiv, 2024–03 (2024)

[21] Del Razo, M.J., Qiu, R., Chen, X., Koes, D.R., Trainor, K., Dill, K.A., Donadio, D.: Sampling conformational space with machine learning. Journal of Chemical Theory and Computation 18(10), 5942–5956 (2022)

[22] Jing, B., Berger, B., Jaakkola, T.: Alphafold meets flow matching for generating protein ensembles. arXiv preprint arXiv:2402.04845 (2024)

[23] Zheng, S., He, J., Liu, C., Shi, Y., Lu, Z., Feng, W., Ju, F., Wang, J., Zhu, J., Min, Y., et al.: Predicting equilibrium distributions for molecular systems with deep learning. Nature Machine Intelligence 6(5), 558–567 (2024)

[24] Song, Y., Sohl-Dickstein, J., Kingma, D.P., Poole, B., Ho, J., Ermon, S.: Scorebased generative modeling through stochastic differential equations. International Conference on Learning Representations (ICLR) (2021)

[25] Kazantsev, A.V., Rambo, R.P., Karimpour, S., SantaLucia, J., Tainer, J.A., Pace, N.R.: Solution structure of RNase P RNA. RNA 17 6, 1159–71 (2011)

[26] Lee, Y.-T., Degenhardt, M.F.S., Skeparnias, I., Degenhardt, H.F., Bhandari, Y.R., Yu, P., Stagno, J.R., Fan, L., Zhang, J., Wang, Y.-X.: The conformational space of RNase p RNA in solution 637(8048), 1244–1251 10.1038/s41586-024-08336-6

[27] Lewis, S., Hempel, T., Jiménez-Luna, J., Gastegger, M., Xie, Y., Foong, A.Y., Satorras, V.G., Abdin, O., Veeling, B.S., Zaporozhets, I., et al.: Scalable emulation of protein equilibrium ensembles with generative deep learning. bioRxiv, 2024–12 (2024)

[28] Naritomi, Y., Fuchigami, S.: Slow dynamics of a protein backbone in molecular dynamics simulation revealed by time-structure based independent component analysis. The Journal of Chemical Physics 139(21) (2013)

[29] Hahn, D.F., Gapsys, V., Groot, B.L., Mobley, D.L., Tresadern, G.: Current state of open source force fields in protein–ligand binding affinity predictions. Journal of Chemical Information and Modeling 64(13), 5063–5076 (2024)

[30] Saldaño, T., Escobedo, N., Marchetti, J., Zea, D.J., Mac Donagh, J., Velez Rueda, A.J., Gonik, E., García Melani, A., Novomisky Nechcoff, J., Salas, M.N., Peters, T., Demitroff, N., Fernandez Alberti, S., Palopoli, N., Fornasari, M.S., Parisi, G.: Impact of protein conformational diversity on AlphaFold predictions. Bioinformatics 38(10), 2742–2748 (2022) 10.1093/bioinformatics/btac202

[31] Jing, B., Erives, E., Pao-Huang, P., Corso, G., Berger, B., Jaakkola, T.: Eigenfold: Generative protein structure prediction with diffusion models. arXiv preprint arXiv:2304.02198 (2023)

[32] Henzl, M.T., Tanner, J.J.: Solution structure of Ca2+-free rat β-parvalbumin (oncomodulin). Protein Science 16(9), 1914–1926 (2007)

[33] Silva, A.M.N., Moniz, T., de Castro, B., Rangel, M.: Human transferrin: An inorganic biochemistry perspective. Coordination Chemistry Reviews 449, 214186 (2021) 10.1016/j.ccr.2021.214186

[34] Halbrooks, P.J., Mason, A.B., Adams, T.E., Briggs, S.K., Everse, S.J.: The oxalate effect on release of iron from human serum transferrin explained. Journal of Molecular Biology 339(1), 217–226 (2004) 10.1016/j.jmb.2004.03.049

[35] Biasini, M., Schmidt, T., Bienert, S., Mariani, V., Studer, G., Haas, J., Johner, N., Schenk, A.D., Philippsen, A., Schwede, T.: OpenStructure: an integrated software framework for computational structural biology. Biological crystallography 69(5), 701–709 (2013)

[36] Buttenschoen, M., Morris, G.M., Deane, C.M.: PoseBusters: AI-based docking methods fail to generate physically valid poses or generalise to novel sequences. Chemical Science 15, 3130–3139 (2024) 10.1039/D3SC04185A

[37] Eddy, S.R.: Accelerated profile HMM searches. PLoS computational biology 7(10), 1002195 (2011)

[38] Suzek, B.E., Wang, Y., Huang, H., McGarvey, P.B., Wu, C.H., Consortium, U.: Uniref clusters: a comprehensive and scalable alternative for improving sequence similarity searches. Bioinformatics 31(6), 926–932 (2015)

[39] Mitchell, A.L., Almeida, A., Beracochea, M., Boland, M., Burgin, J., Cochrane, G., Crusoe, M.R., Kale, V., Potter, S.C., Richardson, L.J., et al.: Mgnify: the microbiome analysis resource in 2020. Nucleic acids research 48(D1), 570–578 (2020)

[40] Steinegger, M., Mirdita, M., Söding, J.: Protein-level assembly increases protein sequence recovery from metagenomic samples manyfold. Nature methods 16(7), 603–606 (2019)

[41] Steinegger, M., Meier, M., Mirdita, M., Vöhringer, H., Haunsberger, S.J., Söding, J.: Hh-suite3 for fast remote homology detection and deep protein annotation. BMC bioinformatics 20, 1–15 (2019)

[42] Wheeler, T.J., Eddy, S.R.: nhmmer: DNA homology search with profile HMMs. Bioinformatics 29(19), 2487–2489 (2013)

[43] Kalvari, I., Nawrocki, E.P., Ontiveros-Palacios, N., Argasinska, J., Lamkiewicz, K., Marz, M., Griffiths-Jones, S., Toffano-Nioche, C., Gautheret, D., Weinberg, Z., et al.: Rfam 14: expanded coverage of metagenomic, viral and microRNA families. Nucleic acids research 49(D1), 192–200 (2021)

[44] Consortium, R.: Rnacentral 2021: secondary structure integration, improved sequence search and new member databases. Nucleic Acids Research 49(D1), 212–220 (2020) 10.1093/nar/gkaa921

[45] Sayers, E.W., Bolton, E.E., Brister, J.R., Canese, K., Chan, J., Comeau, D.C., Farrell, C.M., Feldgarden, M., Fine, A.M., Funk, K., et al.: Database resources of the national center for biotechnology information in 2023. Nucleic acids research 51(D1), 29 (2022)

[46] Steinegger, M., Söding, J.: Clustering huge protein sequence sets in linear time. Nature communications 9(1), 2542 (2018)

[47] Scherer, M.K., Trendelkamp-Schroer, B., Paul, F., Pérez-Hernández, G., Hoffmann, M., Plattner, N., Wehmeyer, C., Prinz, J.-H., Noé, F.: PyEMMA 2: A software package for estimation, validation, and analysis of markov models. Journal of chemical theory and computation 11(11), 5525–5542 (2015)

[48] Kufareva, I., Abagyan, R.: Methods of protein structure comparison. Homology modeling: Methods and protocols, 231–257 (2012)

[49] Mariani, V., Biasini, M., Barbato, A., Schwede, T.: lddt: a local superpositionfree score for comparing protein structures and models using distance difference tests. Bioinformatics 29(21), 2722–2728 (2013)

[50] Zhang, Y., Skolnick, J.: Scoring function for automated assessment of protein structure template quality. Proteins: Structure, Function, and Bioinformatics 57(4), 702–710 (2004)

[51] Lensink, M.F., Wodak, S.J.: Docking, scoring, and affinity prediction in capri. Proteins: Structure, Function, and Bioinformatics 81(12), 2082–2095 (2013)

[52] Basu, S., Wallner, B.: Dockq: a quality measure for protein-protein docking models. PloS one 11(8), 0161879 (2016)

[53] RDKit: Open-source cheminformatics software. https://www.rdkit.org.

[54] Morris, G.M., Huey, R., Lindstrom, W., Sanner, M.F., Belew, R.K., Goodsell, D.S., Olson, A.J.: Autodock4 and autodocktools4: Automated docking with selective receptor flexibility. Journal of Computational Chemistry 30(16), 2785–2791 (2009)

